# Partial duplication of the *CRYBB1-CRYBA4* locus is associated with autosomal dominant congenital cataract

**DOI:** 10.1101/042515

**Authors:** Owen M. Siggs, Shari Javadiyan, Shiwani Sharma, Emmanuelle Souzeau, Karen M. Lower, Deepa A. Taranath, Jo Black, John Pater, John G. Willoughby, Kathryn P. Burdon, Jamie E. Craig

## Abstract

Congenital cataract is a rare but severe paediatric visual impediment, often caused by variants in one of several crystallin genes that produce the bulk of structural proteins in lens. Here we describe a pedigree with autosomal dominant isolated congenital cataract and linkage to the crystallin gene cluster on chromosome 22. No rare single nucleotide variants or short indels were identified by whole-exome sequencing, yet copy number variant analysis revealed a duplication spanning both CRYBB1 and CRYBA4. While the CRYBA4 duplication was complete, the CRYBB1 duplication was not, with the duplicated CRYBB1 product predicted to create a gain of function allele. This association suggests a new genetic mechanism for the development of isolated congenital cataract.

**Grant information:** Supported by the National Health and Medical Research Council

**Conflict of interest:** the authors declare no conflict of interest.

## Introduction

Cataract is an opacification of the crystalline lens and one of the leading causes of blindness worldwide ^1^. Those that occur within the first year of life are categorized as congenital or infantile cataract, with an incidence in the order of 52.8 per 100,000 children ^2^. Around 23% of non-syndromic congenital cataracts are familial ^3^, with around 50% of these associated with a crystallin gene variant ^4^.

Crystallin proteins account for more than 90% of soluble lens protein, and can be divided into α-, β- and γ-crystallin families. With age, the closely related β- and γ-crystallins gradually form insoluble aggregates, which the chaperone-like α-crystallins serve to counteract. Ten human crystallin genes are known to be mutated in congenital cataract, with variants typically thought to reduce crystallin solubility directly (e.g. by creating an insoluble β- or γ-crystallin) or indirectly (e.g. due to loss of α-crystallin chaperone function) ^5^. These genes include both α-crystallins (*CRYAA* and *CRYAB*), two acidic β-crystallins (*CRYBA1* and *CRYBA4*), three basic β-crystallins (*CRYBB1*, *CRYBB2*, and *CRYBB3*), and three γ-crystallins (*CRYGC*, *CRYGD*, and *CRYGS*) ^4^.

The chromosomal arrangement of human crystallin genes reflects their evolutionary history, with major clusters on chromosomes 2 and 22 ^6^. Of a total of eight γ-crystallin genes, six are located on chromosome 2. Similarly, all three basic β-crystallin genes (*CRYBB1*, *CRYBB2*, and *CRYBB3*) are located on chromosome 22, with the acidic β-crystallin *CRYBA4* directly adjacent to *CRYBB1* (but transcribed in the opposite direction). This *CRYBB1-CRYBA4* arrangement is present in organisms as distant as zebrafish, and likely significant for their coordinate regulation. Either gene can lead to congenital cataract when mutated: *CRYBA4* missense variants are known to cause dominant cataract ^7^, while *CRYBB1* variants may be dominant ^8^ or recessive ^9^.

Here we describe an autosomal dominant congenital cataract pedigree associated with a unique duplication of the paired *CRYBB1-CRYBA4* locus. Both genes were found to be duplicated, with a complete duplication of *CRYBA4* and partial duplication of *CRYBB1*.

## Materials & Methods

### Patients

Clinical information from 19 members of pedigree CSA106 (Caucasian) was collected by referring ophthalmologists (see Table 1), with blood samples taken after informed written consent. Of these, 11 had developed bilateral cataracts. The study was approved by the Southern Adelaide Clinical Human Research Ethics Committee.

**Table 1:** Clinical details of CSA106 family members. All affected members had bilateral cataracts. y, years; LE, left eye; RE, right eye; VA, visual acuity; NA, not available.

| ID | Age diagnosed (y) | Exome sequenced | Disease Status | Age of surgery (y) (RE) | Age of surgery (y) (LE) | pre-surgical VA (RE) | pre-surgical VA (LE) |
| --- | --- | --- | --- | --- | --- | --- | --- |
| CSA106.01 | 5 | Yes | Affected | 76 | 76 | NA | NA |
| CSA106.02 | 0 | Yes | Affected | 44 | 44 | NA | NA |
| CSA106.03 | 0 | Yes | Affected | 3 | 22 | NA | NA |
| CSA106.04 | 3 | Yes | Affected | 18 | 19 | NA | NA |
| CSA106.05 | - | Yes | Unaffected | - | - | - | - |
| CSA106.06 | 4 | No | Affected | 7 | 13 | 6/36 | 6/12 |
| CSA106.07 | 0 | No | Affected | NA | NA | 6/15 | 6/15 |
| CSA106.08 | - | Yes | Unaffected | - | - | - | - |
| CSA106.09 | - | Yes | Unaffected | - | - | - | - |
| CSA106.10 | - | No | Unaffected | - | - | - | - |
| CSA106.11 | - | No | Unaffected | - | - | - | - |
| CSA106.12 | NA | No | Affected | NA | NA | NA | NA |
| CSA106.13 | NA | No | Affected | NA | NA | NA | NA |
| CSA106.14 | 10 | Yes | Affected | 51 | 10 | NA | NA |
| CSA106.15 | - | Yes | Unaffected | - | - | - | - |
| CSA106.16 | NA | No | Affected | 24 | NA | NA | NA |
| CSA106.17 | - | Yes | Unaffected | - | - | - | - |
| CSA106.18 | - | No | Unaffected | - | - | - | - |
| CSA106.19 | NA | Yes | Affected | NA | NA | NA | NA |

### Candidate gene panel screening

Individual CSA106.03 (Figure 1A) was screened for variants in 51 known congenital cataract genes using an Ion AmpliSeq custom amplicon panel (Life Technologies). Genomic DNA concentration was measured with a dsDNA high sensitivity Assay Kit on a Qubit fluorometer (Life Technologies). Library preparation (Ion AmpliSeq library kit v2.0) and template preparation (Ion PGM Template OT2 200 Kit) were performed according to the manufacturer’s instructions. The clonally amplified library was enriched on the Ion OneTouch enrichment system and quantified with a Bioanalyzer 2100 using the High Sensitivity DNA Kit (Agilent Technologies). Sequencing was performed on an Ion Torrent Personal Genome Machine (PGM) using The Ion PGM Sequencing 200 Kit v2 and an Ion 318 chip (Life Technologies). Torrent Suite (v3.6) was used to align reads to the hg19 reference genome. The number of mapped reads, percentage of on-target reads, and mean read depth were calculated using the Coverage Analysis plugin (v4.0-r77897), and variants were called using the Variant Caller plugin (v4.0-r76860) with germline algorithm. Ion Reporter v4.0 was used for annotation. Variants were prioritized for further analysis if they were predicted to alter protein coding sense, were rare (MAF<0.001 in Exome Variant Server (EVS) (http://evs.gs.washington.edu/EVS/)), and were absent from unaffected controls.

**Figure 1.**
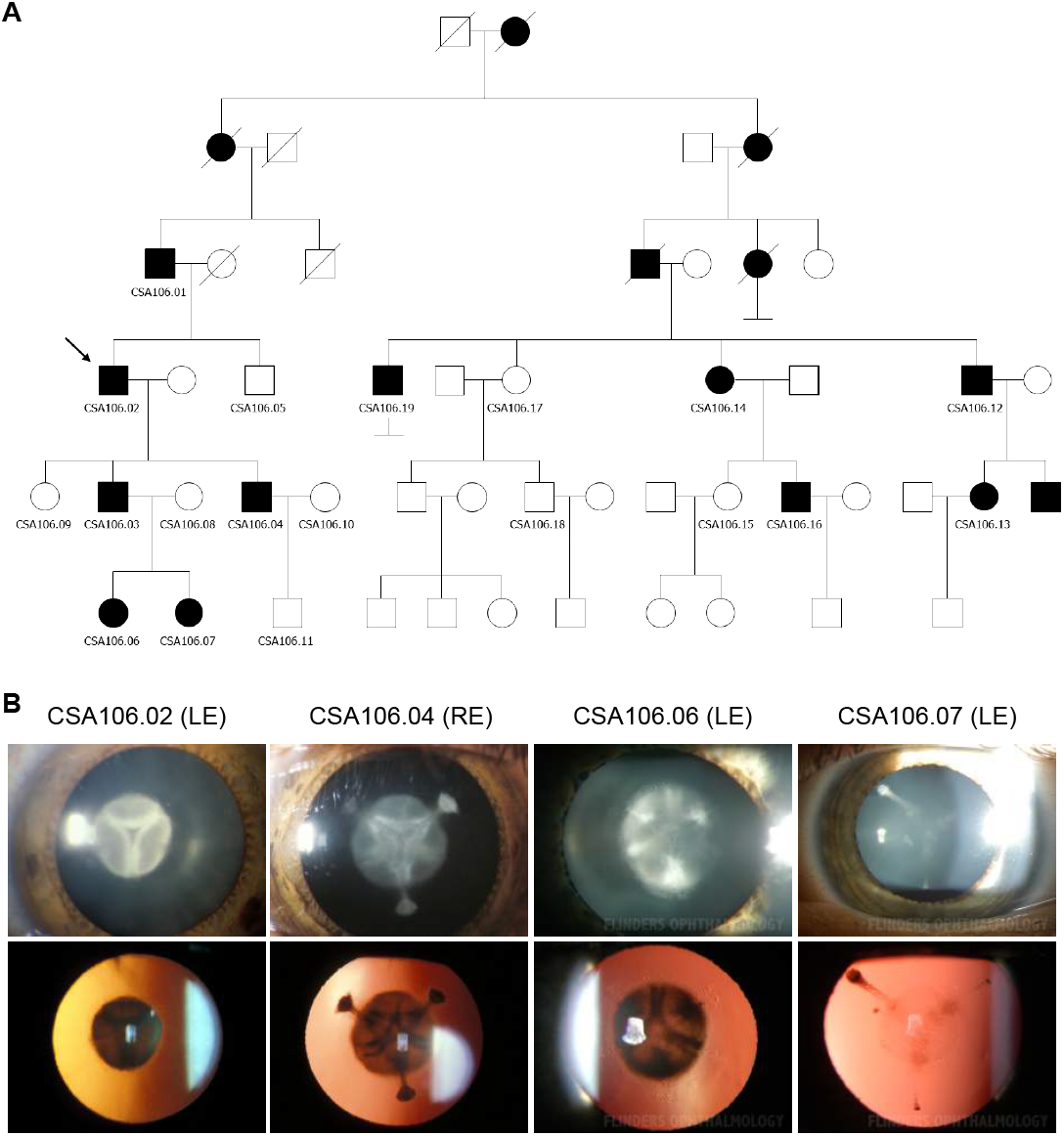
An autosomal dominant congenital cataract pedigree. (A) CSA106 pedigree, indicating affected (black) and unaffected (white) members. Proband (CSA106.02) is indicated by an arrow. (B) Direct illumination (top panels) or retroillumination (bottom panels) of the same cataract from four family members, demonstrating mild to dense fetal nuclear opacification with sutural involvement. LE, left eye; RE, right eye.

### Exome sequencing

A total of 11 members of the CSA106 family (6 affected, 5 unaffected) were subjected to exome sequencing, along with an additional 325 unrelated Australian cases and controls (a mixture of examined normal controls [20], cataract cases [22], keratoconus cases [51], advanced glaucoma cases [195], and primary congenital glaucoma cases [37]). Genomic DNA was extracted from blood samples using a QIAamp Blood Maxi Kit (Qiagen) and subjected to exome capture (Agilent SureSelect v4). Paired-end libraries were then sequenced on an Illumina HiSeq 2000 by an external contractor (Axeq Technologies). Reads were mapped to the human reference genome (hg19) using the Burrows-Wheeler Aligner (v0.7.10), and duplicates were marked and removed using Picard (v1.126). An average of 49,429,126 reads per sample were mapped to capture regions, with a mean read depth of 83.6 and >10X coverage for 97.3% of the capture regions. Variants were called using SAMtools (v1.0) and annotated against RefSeq transcripts using ANNOVAR (2014Nov12) with additional annotations from the NHLBI Exome Sequencing Project (ESP6500SI-V2), 1000 Genomes Project (November 2014 release), ExAC (v0.3.1), and dbSNP138 databases. Variants were filtered by QUAL score (>20), and considered to be potentially pathogenic if predicted to alter protein coding sense (nonsynonymous, stopgain, stoploss, frameshift, essential splice), and were sufficiently rare (ExAC MAF<0.0001). *CRYBB1* variants were annotated according to RefSeq accession numbers NG_009826.1, NM_001887.3 and NP_001878.1, with *CRYBA4* variants based on NG_009825.1, NM_001886.2 and NP_001877.1.

### Linkage analysis

VCF files were converted to MERLIN input format using the vcf2linkdatagen and linkdatagen scripts ^10^. Parametric linkage analysis was then performed using MERLIN (v1.1.2) under a fully penetrant dominant model with a disease frequency of 0.0001.

### Exome CNV analysis

Coverage depth across the critical region was extracted from exome BAM files using SAMtools (v1.3). For copy number variant analysis using CoNIFER (v0.2.2), the same interval was analysed in 343 population-matched control exomes (including 11 from family CSA106) using the following parameters: SVD 5, ZRPKM 1.5.

### qPCR CNV analysis

TaqMan Copy Number Assays for intron 3 (Hs04088405_cn: hg19 chr22:g.27006444) and exon 6 (Hs00054226_cn: hg19 chr22:g.26995522) of *CRYBB1* were ordered from ThermoFisher Scientific. All available CSA106 family members were tested for partial duplication in *CRYBB1* gene using genomic DNA according to manufacturer's protocol. Briefly, each region was amplified in four replicates on a StepOne Plus real-time PCR instrument alongside an endogenous reference gene (TaqMan Copy Number Reference Assay, human, RNase P). CopyCaller 2.0 software (Life Technologies) was used to predict the copy number of the target genomic DNA. Intron 3 was also screened in an additional 46 congenital cataract probands with an unidentified genetic cause (recruited under the same protocol as the CSA106 family).

### Whole-genome sequencing

Genomic DNA from a single affected family member (CSA106.19) was extracted from a blood sample using a QIAamp Blood Maxi Kit (Qiagen). A 150bp paired-end library was generated using the TruSeq Nano kit (v2.5) and sequenced on an Illumina HiSeq X Ten system by an external contractor (Garvan Institute of Medical Research). 148.75Gb of sequence data was generated, with reads mapped to the human reference genome (hg19) using the Burrows-Wheeler Aligner (v0.7.12), and duplicates marked and removed using Picard (v2.6). 988,463,168 reads were mapped to the hg19 reference genome, with a mean read depth of 43.3 and >10X coverage for 96.8% of the genome. Local realignment and base quality recalibration was performed using the Genome Analysis Toolkit (GATK, v3.5). The predicted pathogenic variant identified in this study has been submitted to the ClinVar Database (accession number SCV000484507, https://www.ncbi.nlm.nih.gov/clinvar/).

### Lens protein extraction

Cataractous lens material was collected from the proband during phacoemulsification (CSA106.6, aged 13 years), and stored in balanced salt solution with 2mM EDTA pH 8.0 at −80°C. Normal human lens was obtained from an 18 year-old deceased donor (Eye Bank of South Australia, Flinders Medical Centre), collection of which was approved by Southern Adelaide Clinical Human Research Ethics Committee. Lenses were homogenized in 2 mL of extraction buffer containing 50 mM imidazole (pH 7), 50 mM NaCl, 2 mM 6-aminohexanoic acid, 1 mM EDTA, and protease inhibitor cocktail (Roche Diagnostics), and ultracentrifuged at 150,000×*g* for 30 min at 4°C. The soluble fraction was acetone precipitated according to the Thermos Scientific protocol (Thermo Fisher). The insoluble fraction was dissolved in buffer containing 6 M urea, 2% dichloro-diphenyl-trichloroethane (DTT), 2% 3-[C3-cholamidoproyl] dimethyl-ammonio-1-propansulfonat (CHAPS) and 0.1% sodium dodecyl sulfate (SDS). The EZQ Protein Quantitation method (Life Technologies) was used to determine protein concentration.

### Denaturing gel electrophoresis and Western blotting

Twenty micrograms of total soluble protein from each lens was size fractionated by SDS-PAGE using a 12% polyacrylamide gel. The precision plus protein standards (BioRad) were used for size comparison. Fractionated proteins were transferred onto Hybond-C Extra nitrocellulose (GE Healthcare), and after blocking in 5% (wt/vol) milk in TBS-Tween (Tris Buffered Saline and 1% Tween-20) was incubated with a mouse monoclonal anti-CRYBB1 primary antibody (1:400, Sigma-Aldrich, WH0001414M3). After washing the membrane was incubated with horseradish peroxidase (HRP)-conjugated goat anti-mouse IgG (1:1000, Jackson ImmunoResearch), and following another wash was treated with Clarity Western ECL Blotting Substrate (BioRad) or Amersham ECL Prime (GE Healthcare) and imaged using an ImageQuant LAS 4000 Imager (GE Healthcare). The same membrane was stripped at 50°C in stripping buffer (100mM β-mercaptoethanol, 2% SDS and 62.5 mM Tris-HCl [pH 7]) then reprobed with polyclonal rabbit anti-CRYBA4 (1:200, Abcam, ab130680) or polyclonal sheep anti-CRYΑA (1:1000; Flinders University Antibody Production Facility) primary antibodies, followed by HRP-conjugated goat anti-rabbit IgG or anti-sheep IgG secondary antibodies (1:1000, Jackson ImmunoResearch).

## Results

### Autosomal dominant congenital cataract

We identified a 6-generation autosomal dominant congenital cataract pedigree (Figure 1A), with multiple affected members diagnosed with nonsyndromic bilateral cataracts between birth and 10 years of age (Table 1). Slit lamp imaging of the lens demonstrated mild to dense fetal nuclear opacification with anterior and posterior sutural involvement (Figure 1B). A custom amplicon sequencing panel, designed to sequence 51 known congenital cataract genes (Supplementary Table 1), was first used to screen an affected family member (CSA106.03). A total of 807,055 reads were mapped to the reference genome, covering 93.21% of target gene bases to at least 20X depth (average read depth of 623.5 across 1216 amplicons). A total of 172 variants were identified: all 172 either had a mean allele frequency (MAF) of greater than 0.001, were present in control samples, or were not predicted to alter protein sequence.

### Linkage to chromosome 22

Given the absence of a candidate variant in a known congenital cataract gene that affects function, we sequenced the exomes of 11 family members (6 affected, 5 unaffected). Parametric linkage analysis under a rare dominant inheritance model revealed a peak LOD score of 3.3 on chromosome 22 (Figure 2A,B). Haplotype phasing indicated that rs2236005 (hg19 chr22:g.26422980A>G) and rs2347790 (hg19 chr22:g.29414001A>G) were the boundaries of the 3 Mbp critical region (Figure 2C), within which lay 17 protein-coding genes (plus 9 ncRNA and 7 pseudogenes) including the known congenital cataract genes *CRYBB1* and *CRYBA4* (Figure 2D). All 17 protein-coding gene exons were covered by whole-exome capture sequencing, with *CRYBB1* and *CRYBA4* also having been previously covered by candidate gene amplicon sequencing. We detected two single-nucleotide variants that were shared between affected family members, absent from unaffected family members, and that altered coding sense (Figure 2E). Both variants had a MAF at least an order of magnitude greater than the estimated population incidence of congenital cataract (~0.000528), and therefore were considered extremely unlikely candidates. Synonymous variants in *CRYBB1* or *CRYBA4* – recently identified as a possible cause of crystallin misfolding ^11^ – were also not shared between affected members.

**Figure 2.**
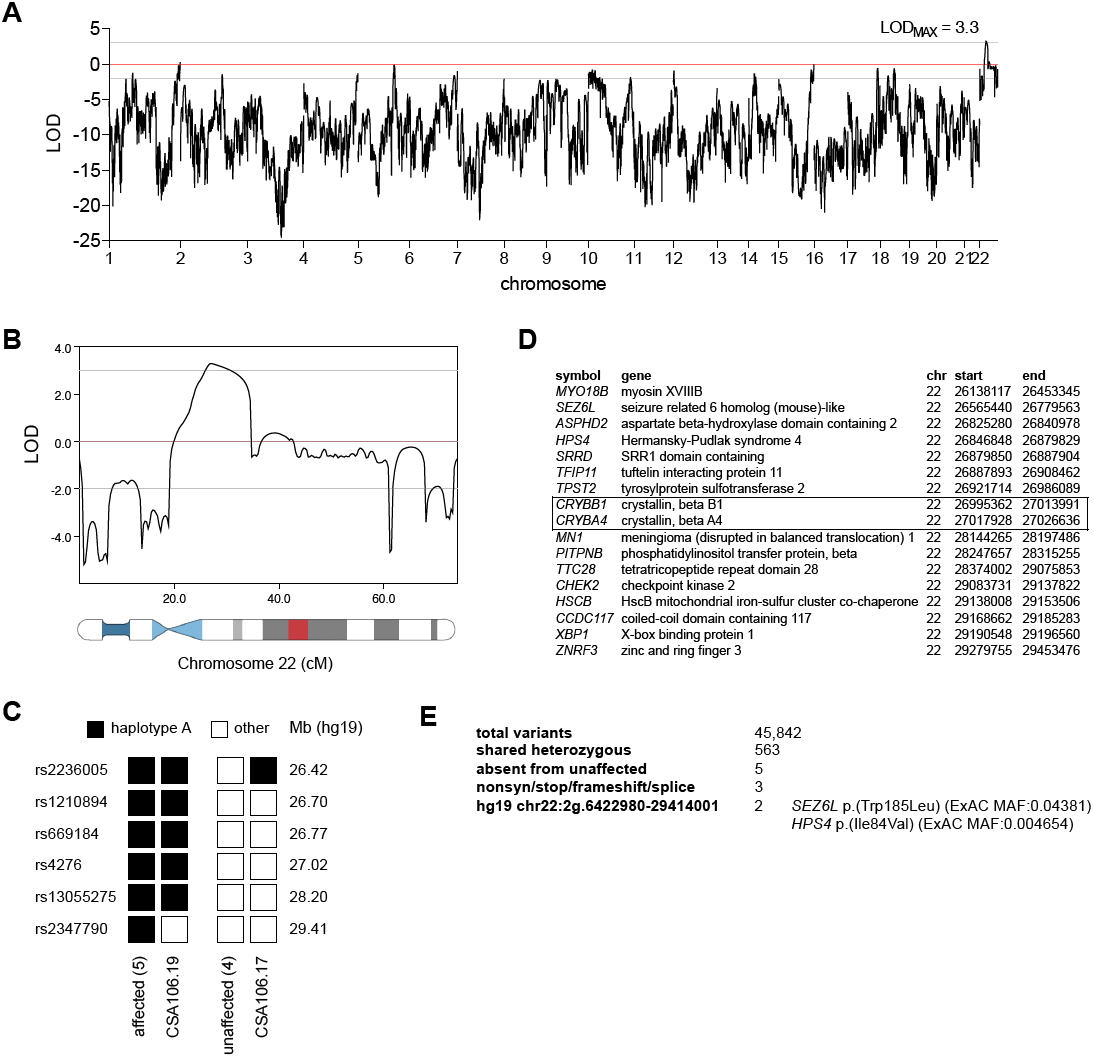
Linkage of the cataract phenotype to proximal chromosome 22q. (A) Genome-wide LOD scores under a fully penetrant dominant inheritance model. (B) LOD scores across chromosome 22. (C) Haplotype analysis and critical recombinants across the interval. (D) List of protein-coding genes within the defined linkage interval, with previously known congenital cataract genes highlighted. Start and end coordinates refer to the hg19 reference sequence. (E) Filtering strategy to identify shared heterozygous coding variants within linkage interval.

### Partial duplication of the *CRYBB1-CRYBA4* locus

We next investigated coverage depth across the linkage interval using two methods. Both CoNIFER (Figure 3A) and SAMtools (Figure 3B) revealed an increased coverage depth (and by inference increased copy number) at the *CRYBB1*-*CRYBA4* locus of all affected individuals. Based on coverage of protein-coding exons across the locus, this CNV spanned between a minimum of 28.8 kbp (hg19 chr22:g.26997843_27026636) and a maximum of 1.15 Mbp (hg19 chr22:g.26995638_28146902), the smaller of which encompassed only two protein-coding genes (*CRYBB1* and *CRYBA4*), and the larger of which also included a long non-coding RNA gene (*MIAT*). This CNV (hg19 chr22:g.(26995638_26997843)_(27026636_28146902)dup) was present in all six affected family members, absent from all five unaffected family members, and was absent from a further 325 unrelated Australian exomes sequenced contemporaneously. Mean coverage depth analysis revealed that while all five captured exons of *CRYBA4* appeared to be duplicated (representing the final five of six total exons), only the first five exons of *CRYBB1* (of a total of six) appeared to have been duplicated (Figure 3B). Duplication of *CRYBB1* intron 3 was confirmed by qPCR in all affected family members, with a similar assay showing that distal exon 6 was not duplicated (Figure 3C,D). *CRYBB1* and *CRYBA4* variation was also manually inspected via Integrative Genomics Viewer, given that a variant in one *CRYBB1-CRYBA4* locus of a total of three (two alleles plus the duplicated allele) may not be called as heterozygous by variant-calling algorithms. We also screened an additional 46 unsolved congenital cataract cases for *CRYBB1* duplications, yet did not identify any further duplications (Figure 3E).

**Figure 3.**
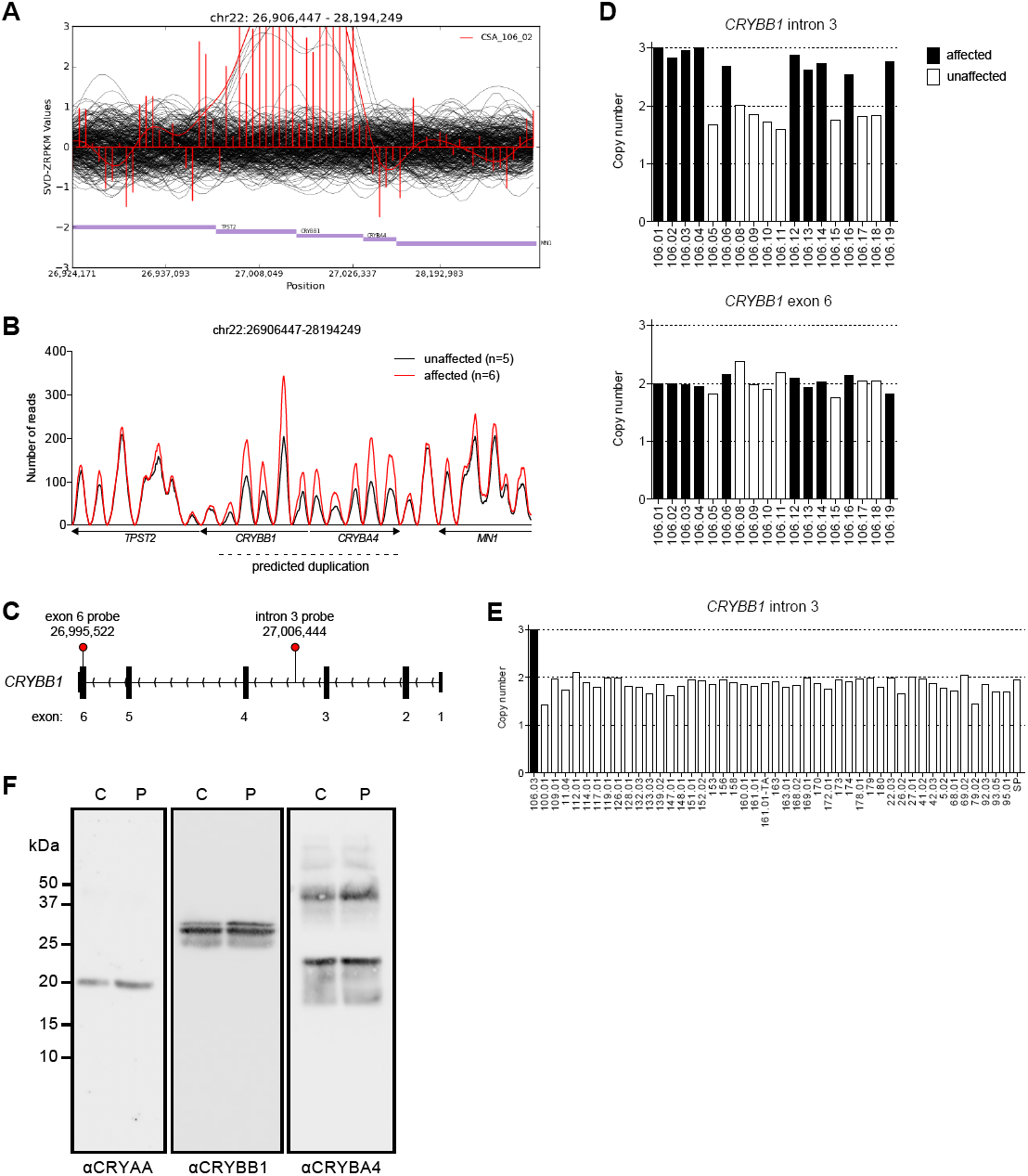
Partial duplication of the CRYBB1-CRYBA4 locus. (A) CoNIFER output indicating a copy number gain within the captured portion of the linkage interval. (B) Average read depth of affected (red) and unaffected (black) family members across the same interval shown in (A). Each peak represents a region covered by exome capture and sequencing, typically an exon. All five coding exons of both *CRYBB1* and *CRYBA4* were covered, with an additional region captured and sequenced for *CRYBB1*. The final peak at the *CRYBB1* locus (i.e. furthest to left) represents the sixth and final exon. (C) Exon structure of *CRYBB1* and location of qPCR probes used for copy number variant analysis. (D) qPCR validation of a duplicated (intron 3) and non-duplicated (exon 6) region of *CRYBB1*. Bars indicate copy number (CN) normalized to an internal control probe. (E) *CRYBB1* duplication screening in unsolved congenital cataract cases. (F) Crystallin protein content in cataractous lens extracts. Control [C] and patient [P] lens extracts were subjected to Western blotting with the indicated antibodies. CRYAA served as a loading control, with the same blot probed sequentially with all three antibodies.

To examine the effects of the *CRYBB1-CRYBA4* duplication on protein expression, we prepared protein from the cataractous left lens of CSA106.06 (removed during phacoemulsifcation) and from a non-cataractous control lens. An anti-CRYAA Western blot indicated equivalent loading between the cataract and control samples, and reprobing the same blot with anti-CRYBA4 and anti-CRYBB1 blots revealed bands of the appropriate size (22kDa and 28kDa monomers, respectively) and similar density in both samples (Figure 3F). An additional band was detected with the anti-CRYBA4 antibody corresponding to a CRYBA4 dimer (44kDa), although no other bands were apparent. We did not detect any additional anti-CRYBB1-reactive bands in the cataract sample in soluble fractions (Figure 3F), or insoluble fractions (data not shown), despite using an antibody raised against a peptide (NP_001878, p.37_138) that is not encoded by exon 6.

### Tandem duplication of the *CRYBB1-CRYBA4* locus

In order to define the nature of the duplication and the associated breakpoints, we performed whole-genome sequencing on genomic DNA from an affected family member (CSA106.19). Read depth analysis confirmed the presence of a 78,928bp duplication at the *CRYBB1-CRYBA4* locus (hg19 chr22:g.26995597_27074524dup), the boundaries of which occurred immediately proximal to chr22:26995597, and distal to chr22:27074524 (Figure 4A). This duplication encompassed the entirety of *CRYBA4* and the long non-coding RNA gene *MIAT*, but only partially involved *CRYBB1*. Importantly, the proximal duplication boundary occurred within exon 6 of CRYBB1, just 75bp distal to the site of the exon 6 qPCR probe used in Figure 3D. This also explains why a duplicated exon 6 was not detected by whole-exome sequencing, as there would be insufficient complementarity to the exon 6 capture probe.

**Figure 4.**
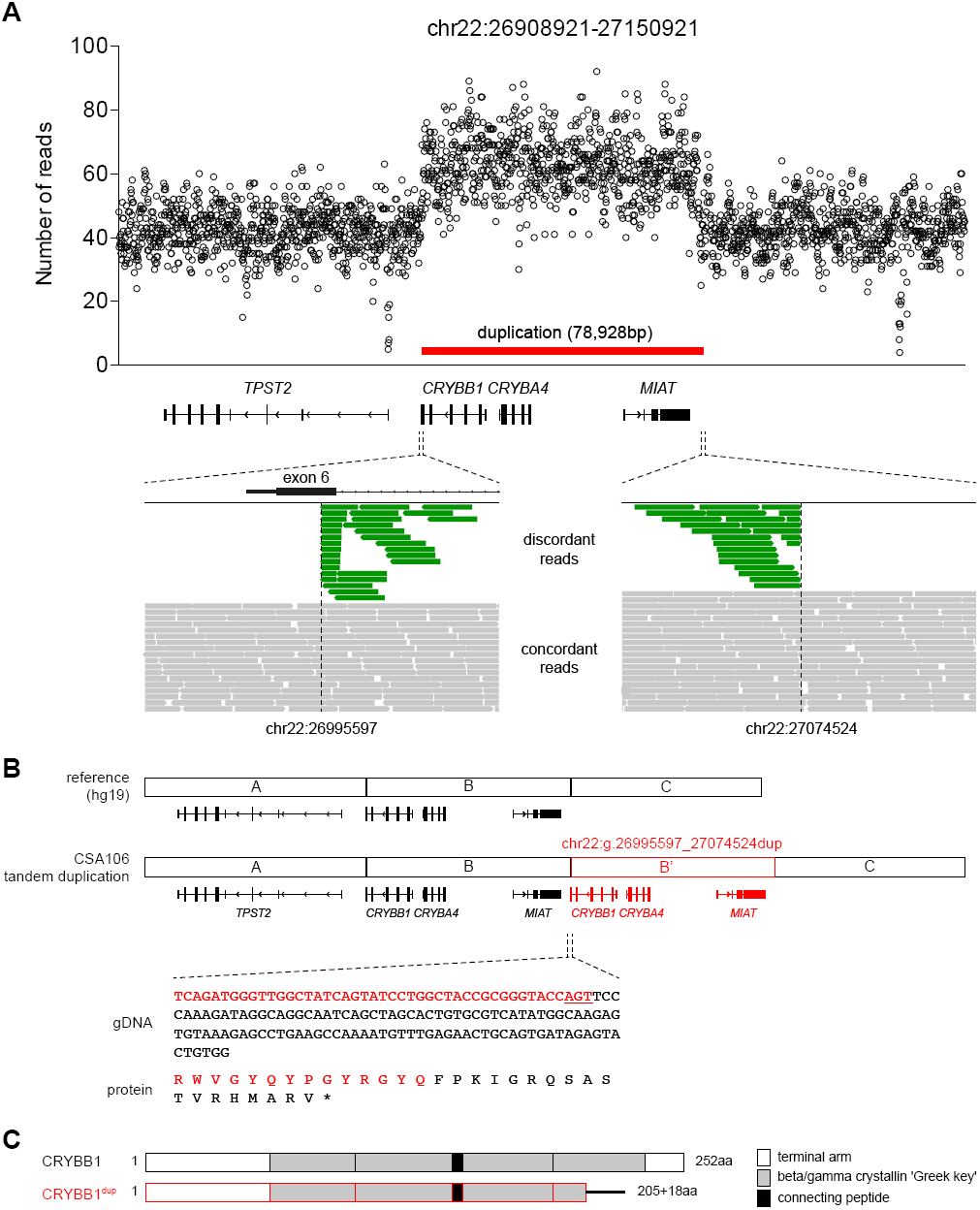
A direct tandem duplication at the *CRYBB1-CRYBA4* locus resulting in partial duplication of *CRYBB1*. (A) Whole-genome sequencing depth across the *CRYBB1-CRYBA4* locus of an affected family member (CSA106.19). Relative location of local transcripts, concordant (grey) and discordant (green) paired reads, and duplication boundaries are indicated below. (B) Depiction of the CSA106 tandem duplication, showing the sequence of a 150bp split read spanning the duplication breakpoint (gDNA) and its predicted translation product (protein). Red gDNA sequence aligns to exon 6 of the duplicated *CRYBB1* gene, with black sequence aligning to a region distal to the long non-coding RNA, *MIAT*. A 3bp region of microhomology is underlined. Predicted translation of the mate pair sequence reveals 14 amino acids from the C-terminus of CRYBB1 (red), followed by an 18 amino acid read-through product and premature termination codon (black). (C) Domain structure of full-length CRYBB1 protein, and hybrid CRYBB1 protein produced as a consequence of the CSA106 tandem duplication (CRYBB1^dup^).

The arrangement of multiple discordant mate pairs within the duplicated region (Figure 4A) suggested a direct tandem duplication ^12^. This was confirmed by BLAST alignments of split reads spanning the duplication breakpoint, which revealed a proximal breakpoint joining exon 6 of *CRYBB1* to intergenic sequence distal to *MIAT* (Figure 4B). No insertions, deletions, or other rearrangements were evident at either proximal or distal breakpoints, although there was 3bp of microhomology at the adjoining sequences of the proximal breakpoint (Figure 4B, underlined). A predicted consequence of this proximal breakpoint was the creation of a hybrid *CRYBB1* gene. This gene encoded the expected CRYBB1 amino acid sequence up to and including amino acid 205 (a glutamine residue). Beyond this point (corresponding to the proximal breakpoint) the final 47 C-terminal amino acids, including the majority of the fourth and final 'Greek key' crystallin domain, are predicted to be replaced by 18 unrelated amino acids encoded by intergenic sequence (Figure 4B, C). If transcribed, this product would also be predicted to avoid nonsense-mediated decay, as the premature termination signal occurs in the final exon.

## Discussion

Here we describe an autosomal dominant congenital cataract pedigree with unambiguous linkage to chromosome 22. Despite the absence of a candidate single nucleotide variant in the coding regions of genes within the linked interval, we identified a partial direct tandem duplication of the *CRYBB1-CRYBA4-MIAT* locus. Although we have not definitively ascribed pathogenicity to this duplication, there are precedents for both *CRYBB1* and *CRYBA4* causing dominant congenital cataract.

Variants in *CRYBA4*, for example, have been described in autosomal dominant congenital cataract ^7,13^. Both reports describe missense variants (c.225G>T (p.(Gly64Trp)), c.242T>C (p.(Leu69Pro)), c.317T>C (p.(Phe94Ser))) which presumably promote cataract formation by creating a less soluble protein. Yet the *CRYBA4* duplication described here covered the complete gene, did not contain any missense variants, and did not lead to any obvious change in protein expression.

On the other hand, *CRYBB1* variants appear to have two distinct pathways to cataractogenesis. The recessive *CRYBB1* variants c.169delG (p.(Gly57Glyfs*107)) or c.2T>A (p.(Met1Lys)) are not expected to be expressed at all, and presumably cause cataracts by completely removing an important structural component ^9,14^. Conversely the dominant alleles such as c.658G>T (p.(Gly220*)) ^8^, c.737C>T (p.(Gln223*)) ^15^ and c.827T>C (p.(*253Arg)) ^16^ are predicted to cause cataract by disrupting the coding sequence of the final exon (exon 6), and creating a protein with reduced solubility. The hybrid *CRYBB1* gene created as a consequence of the duplication described here also disrupts the coding sequence of exon 6, and is associated with an autosomal dominant pattern of inheritance. We therefore consider it most likely that the *CRYBB1* duplication product is the disease-causing agent, despite the fact that we did not detect an appropriately-sized band by Western blotting.

The absence of an additional CRYBB1 band on Western blot may still be consistent with a gain-of-function mechanism: for example, the duplicated CRYBB1 product may have only been transiently expressed, or has a shorter half-life than full-length CRYBB1. Conversely it may not have been synthesised at all, and played no role in cataractogenesis in this family. Given this possibility, we have not excluded the possibility that *CRYBA4* triplosensitivity was responsible for the cataractogenesis, which could conceivably alter the stoichiometry of crystallin subunits at a critical stage of lens development. It is also possible that local transcription is altered in the context of a tandem duplication, again raising implications for crystallin stoichiometry. Duplication of the *MIAT* long noncoding RNA seems the least likely explanation for cataractogenesis in this family, given the large body of evidence for *CRYBB1* and *CRYBA4* in congenital cataract.

More broadly, copy number variation is largely overlooked in many whole-exome and whole-genome studies, perhaps due in part to the limited predicted contribution of CNVs to common disease ^17^. In cases where CNVs do play a role it is almost always deletions that are responsible, either in *trans* with a second deleterious allele, or by causing haploinsufficiency on their own. Increases in copy number are far less common in a disease setting, and when they do occur, they commonly involve complete genes. Ocular disease is no exception, with duplication or triplication of *TBK1* in normal tension glaucoma being one example ^18,19^, and a complex *NHS* triplication in the congenital cataract-associated Nance–Horan syndrome being another ^20^. *TBK1* CNVs associated with glaucoma cover the entire locus, so a mechanistic explanation has not been immediately obvious. In the case of the Nance–Horan syndrome triplication, disruption of *NHS* transcription is thought to explain the phenotype, which is consistent with the loss-of-function mechanism of other *NHS* variants ^20^. In a third example, both deletions and duplications of the same gene (*FOXC1*) have been associated with anterior segment dysgenesis ^21^.

Other diseases can be caused by partial gene duplication ^22^, including ~7% of cases of the X-linked Duchenne and Becker muscular dystrophies (DMD/BMD) ^23^. In the case of DMD these variants invariably cause a frameshift, whereas in BMD the reading frame is maintained ^22^. In both cases the predicted result is a loss or reduction in protein function.

Duplication has been integral to the diversification of the crystallin gene family. In the family presented here, we show that crystallin duplication can also be associated with congenital cataract. This represents a previously undescribed genetic mechanism for the development of isolated congenital cataract, with implications for other inherited diseases that appear refractory to whole-exome or whole-genome sequencing.

## Acknowledgements

We thank Angela Chappell for cataract photography. Supported by the National Health and Medical Research Council (NHMRC) grant 1023911. KPB is funded by an NHMRC Senior Research Fellowship, and JEC by an NHMRC Practitioner Fellowship.

